# TopHat-Recondition: A post-processor for TopHat unmapped reads

**DOI:** 10.1101/033530

**Authors:** Christian Brueffer, Lao H Saal

## Abstract

**Summary:** TopHat is a popular spliced junction mapper for RNA sequencing data, and writes files in the BAM format – the binary version of the Sequence Alignment/Map (SAM) format. BAM is the standard exchange format for aligned sequencing reads, thus correct format implementation is paramount for software interoperability and correct analysis. However, TopHat writes its unmapped reads in a way that is not compatible with other software that implements the SAM/BAM format. We have developed TopHat-Recondition, a post-processor for TopHat unmapped reads that restores read information in the proper format. TopHat-Recondition thus enables downstream software to process the plethora of BAM files written by TopHat.

**Availability and implementation:** TopHat-Recondition is implemented in Python using the Pysam library and is freely available under a 2-clause BSD license on GitHub: https://github.com/cbrueffer/tophat-recondition.

## Introduction

RNA sequencing (RNA-seq) has become as a cornerstone of genomics research. TopHat and TopHat2 ([Trapnell et al., 2009, Kim et al., 2013]; jointly referred to as TopHat from here on) is a highly-cited spliced read mapper for RNA-seq data that is used in many large-scale studies around the world, for example in breast cancer ([Saal et al., 2015]). A search for the term “TopHat” in the NCBI Gene Expression Omnibus (GEO) and the European Nucleotide Archive (ENA) yields 267 and 150 datasets using TopHat, respectively, with the true number being likely much higher.

TopHat writes read data in the BAM format – the binary version of the Sequence Alignment/Map (SAM) format ([Li et al., 2009]), but unlike other read mappers, it writes separate files for reads it could map to the reference genome `(accepted_hits.bam)` and reads it could not map `(unmapped.bam)`. Although many analyses focus on mapped reads alone, often it is necessary to consider unmapped reads, for example to perform quality assurance, to deposit the data in online archives, or to analyze the unmapped reads themselves.

However, all released versions of TopHat to date (version ≤ 2.1.0) generate `unmapped.bam` files that are incompatible with common downstream software, *e.g*., the Picard suite (http://broadinstitute.github.io/picard), SAMtools ([Li et al., 2009]), or the Genome Analysis Toolkit (GATK) ([McKenna et al., 2010]). Even if the problems leading to the incompatibility are corrected in future versions of TopHat, an immense amount of data has already been aligned with affected versions and would need to be realigned, and potentially reanalyzed. TopHat-Recondition is a post-processor for TopHat unmapped reads that corrects the compatibility problems, and restores the ability to process BAM files containing unmapped reads.

## TopHat SAM/BAM-Format Compatibility

TopHat’s `unmapped.bam` incompatibility with other tools has three origins: software bugs resulting in violations of the SAM/BAM specification (https://samtools.github.io/hts-specs/SAMv1.pdf), divergences from the specification’s recommended practices, and different interpretation of acceptable values for some of the file format’s fields between software.

Two TopHat issues impair compatibility: First, all unmapped read-pairs lack the 0x8 bit (next segment in the template unmapped) in their `FLAGS` field. This leads to downstream software incorrectly assuming the reads to be mapped. Second, for unmapped reads where the `FLAGS` field declares the paired read to be mapped, this mapped paired read may be missing from the sequence files. This makes the unmapped read’s fields invalid and can lead to software searching for, and failing to find the paired read.

The SAM/BAM specification contains a section on recommended practices for implementing the format. For read-pairs with one mapped and one unmapped read, TopHat does not follow the recommendations that `RNAME` and POS of the unmapped read should have the same field values as the mapped read. Additionally we found that setting `RNEXT` to the mapped read’s `RNEXT` value, and `PNEXT` to 0 improves compatibility.

Lastly, there are differing interpretations of which field values are acceptable in certain conditions between software packages. For example, the valid range of values for the BAM mapping quality (`MAPQ`) is 0255. For unmapped reads, `TopHat` always sets the `MAPQ` value of unmapped reads to 255, and BWA ([Li and Durbin, 2009]) sets the value to greater than 0 in certain conditions, while the Picard suite asserts that this value be 0 and returns an error when encountering such a read, which can confuse users.

Some BAM-processing software, *e.g*., Picard and GATK can be configured to accept reads that do not conform to its expectations by ignoring errors, thus allowing processing to succeed. However, the resulting BAM files remain non-compliant to the specification which can lead to issues in later analysis steps that are difficult to debug.

## Implementation

TopHat-Recondition is implemented in Python using the Pysam library (https://github.com/pysam-developers/pysam) and requires Python 2.6 or higher. The simplified workflow of the software is shown in figure 1. First, the `unmapped.bam` file is loaded into memory, both for performance reasons and to enable random access to the unmapped reads. In the first pass over the unmapped reads the /1 and /2 suffixes are removed from read names (only `TopHat` prior to version 2.0.7), `MAPQ` is set to 0, missing 0x8 flags are added to unmapped read-pairs, and the reads are indexed by their read names (`QNAME`). In the second pass all unmapped reads with mapped mate are recorded to enable detection of missing mapped mates. The `accepted_hits.bam` file is read sequentially to obtain information to correct unmapped reads with mapped mate; the previously built index is used to quickly access the unmapped mate of the current mapped read. The mate-related bits (0x1, 0x2, 0x8, 0x20, 0x40, 0x80) in the `FLAGS` field of unmapped reads for which the mapped paired read could not be found are unset, effectively making them unpaired. Additionally, the `RNAME, RNEXT, PNEXT` and `POS` fields are modified as described above. The corrected unmapped reads are written as `unmapped_fixup.bam` in the specified directory (by default the input BAM file directory), along with a log file detailing the performed modifications. TopHat-Recondition can process a library with 50 million reads in ten minutes on a standard PC, with the disk read performance being the limiting factor.

**Figure 1:**
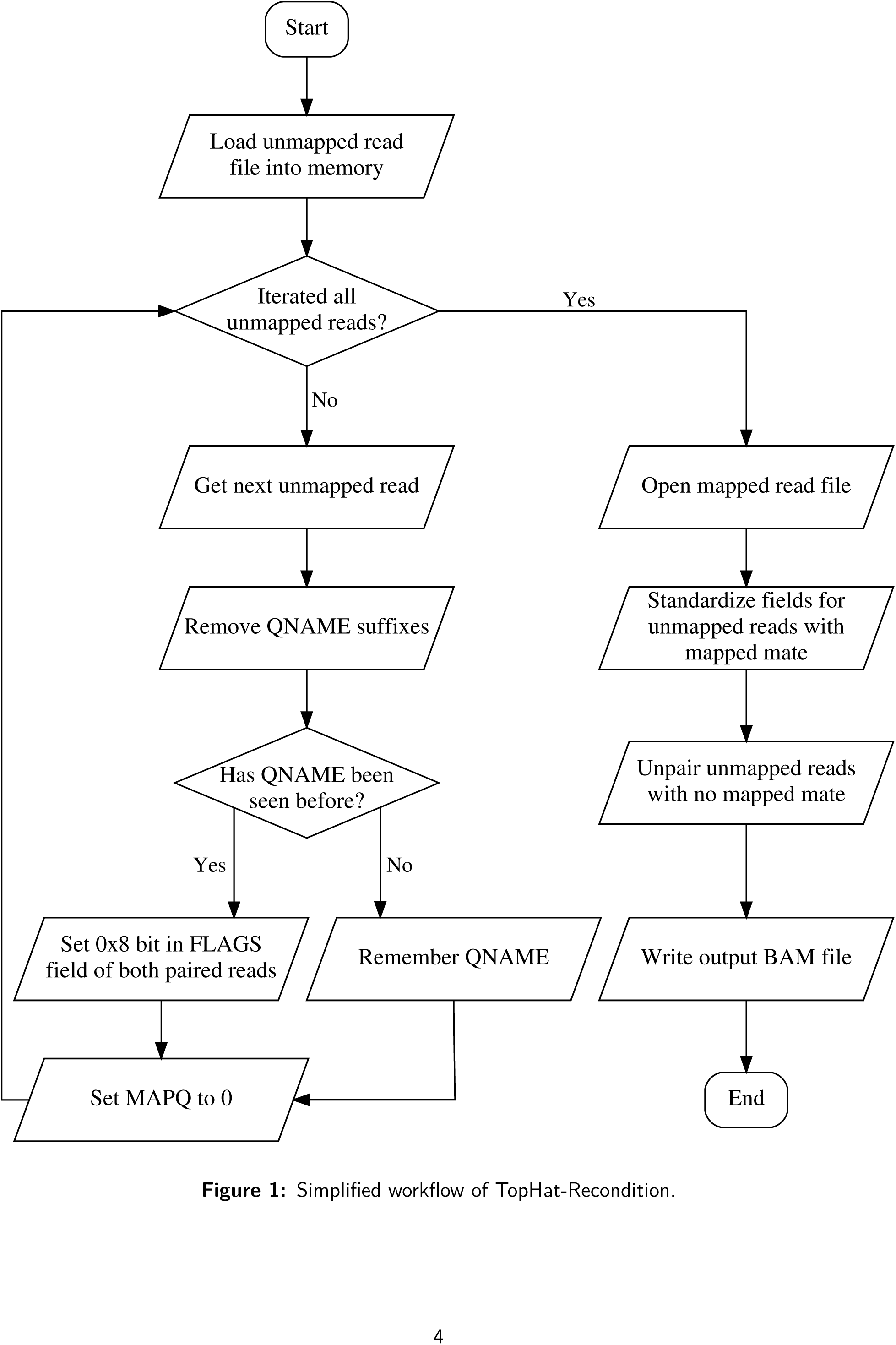
Simplified workflow of TopHat-Recondition.

## Conclusion

TopHat-Recondition enables easy and fast post-processing for TopHat unmapped reads. The tool can be used to process TopHat-written unmapped reads to make them compatible with downstream tools such as samtools, the Picard suite and GATK, which is currently not possible with the stock unmapped reads. This will increase the utility of the immense amount of RNA-seq data that has been analyzed by TopHat.

## Acknowledgements

The authors would like to thank the TopHat developers for permission to use the name TopHat as part of TopHat-Recondition, and Christof Winter for fruitful discussions about the SAM/BAM file format.

